# Evidences of an inhibitory effect and other functions of human P53 in *Escherichia coli*

**DOI:** 10.1101/2024.08.06.606936

**Authors:** Sihem Ben Abid, Ines Yacoubi, Lamia Djemal, Mosbeh Dardouri, Ali Gargouri

## Abstract

The human tumor suppressor P53 has so far been shown to inhibit growth in the yeast model as well as in other eukaryotic contexts. Despite a considerable number of sudies involving p53 in bacteria, the question of effects on cell growth and viability has never been explored. In this work, we report similar negative effect on cell viability of the protein expressed in *Escherichia coli* strain BL21(DE3). This inhibition still needs to be characterized in lights of the distinction between yeast and other organisms with different P53-caused deaths and pathways. Primary tests leaned towards an active p53 in this bacterial context, both under normal and stresseful conditions. Special effects were noticed either with phage infection or antibiotic treatment, using a GST-p53 fusion form. These results open the way for further investigations involving P53 in prokaryote systems.

## Introduction

Human tumor suppressor P53 is a transcription factor that plays key roles in cellular mechanisms such as cell cycle control and arrest, decision and execution of cell death and viability, metabolism and cellular energy **[1]**. The protein has been expressed in many heterologous prokaryotic **[2, 3]** and eukaryotic systems : Leishmania **[4]**, baculovirus infected insect cells **[5]**, etc.

Among the functionalities of ectopic p53 found in some organisms is growth inhibition. This is proven in eukaryotes such as yeast **[6]** ; where apoptotic death is observed when P53 is overexpressed **[7–9]**.

On the other hand, *E. coli* and prokaryotes have been explored for cell death mechanisms. Prokaryotes can induce cell death under certain conditions such as thymine starvation, known as thymine-less death (TLD) as reported by **[10–13]**. This TLD process has been suggested to be associated with the presence of ROS (Reactive Oxygen Species) in bacteria **[14]**.

The p53 gene has been widely expressed in prokaryotes, mainly in *E. coli* with the aim of using it for protein purification or for some protein interactions studies. However, to our knowledge, none of these studies have addressed the possible effects of this tumor suppressor on the growth and physiological behavior of the bacterium. Nonetheless bacterially expressed P53 protein showed functionality in a cell-free apoptosis test assay and was able to interact with mitochondria, induce mitochondrial outer membrane permeabilization (MOMP) and associated apoptotic events **[3]**. When delivered into a solid tumor mouse model via a tumor-specific targeting mechanism, P53-expressing bacteria showed an antitumor effect **[15]**.

In this work, we investigated the question wether recombinant P53 can show any effect on cell physiology and in particular on cell growth of *E. coli* as with other organisms such as yeast. We tried different temperatures for the culture of bacteria expressing P53, mainly 37 °C and 30 °C. Temperatures of 25-30°C were tested based on the fact that low temperatures generally disfavor the aggregation and insolubility of recombinant protein in cells **[16]**, P53 being very prone to aggregation and insolubility as demonstrated by many researches **[17]**. Low Temperatures were investigated in order to avoid this eventual insolubility and to favor any possible protein functionality. And we collected some relevant observations related to this expression of human p53 in bacteria indicating some functionality.

## Material and methods

### Esherichia coli strains

Recombinants BL21 (DE3) strains harboring either empty pGEX4T3 or recombinant pGEX4T3 expressing P53 as in **[2]**.

DH5α, TG1, XL1Blue and TOP10F’. All strains were transformed with the pGEX4T3 vector expressing GST-P53 fusion.

The strain ER2738 from NEB (Phage Display libraries kit) was used to amplify phages as in **[2]**.

### Phages

Ph6, displaying NG7 peptide NPNSAQG, previously isolated from phage display screening against P53 ; Ph17, an empty non recombinant phage and phage control M13mp18 **[2]**.

### Expression of recombinant P53 in bacterial strains

P53 expressing strains (1: BL21 (DE3), 2 : DH5α, 3 : TG1, 4 : TOP10 and 5 : XL1Blue) were cultured at 37°C until OD_600_ of 0.6-0.8 then induced with 0.5 mM IPTG for 3h at 30°C. Cells were sonicated in presence of MOPS buffer supplemented with PMSF 1mM and Lysozyme 0.1 mg/ml. Lysis supernatants were migrated in SDS-gels and used for an immunoblot to detect P53 protein with p53 antibody (Pab 1801): sc-98 (Santa Cruz).

### Kinetics of Bl21 GST-P53 on liquid LB medium

Three strains (non recombinant BL21, BL21/GST and BL21/GST-P53) were cultured in the presence of 0.2 mM IPTG, in parallel with an uninduced BL21/GST-P53 strain, at two temperatures 25°C and 37°C. The non-recombinant strain was cultured on LB while the other two were cultured on LBA (ampicillin). Culture duration: ∼14-15h ; and it was even continued overnight. Sampling and monitoring of OD600 were performed every ∼1h.

### Kinetics with ampicillin supplementation in induced culture medium

Three bacteria strains : untransformed BL21(DE3), BL21(DE3) expressing respectively GST and the fusion GST-P53 are cultured at 25°C. Both transformants were induced with 0.5 mM IPTG from the beginning. Ampicillin absent from the media was introduced (100 µg/ml) - not at the beginning but - in the middle of the exponential phase of the two recombinant strains.

### Phage infection assay performed on strain Bl21/GST-P53

BL21/GST-P53 strain culture induced with IPTG in early exponential phase at 30°C was divided into distinct batches infected separately with three phages : Ph6 (displaying the NG7 peptide NPNSAQG) a P53-specific phage, Ph17 a non-recombinant empty phage, both isolated from phage display liraries in a previous work **[2]** and M13 : M13mp18. Phages used for each infection were simultaneously from amplified and non-amplified solutions. With a control non treated (NT). Infection was let overnight at 30°C 150rpm, tubes were stored at 4°C and cultures were migrated on SDS-denatured gel and immunoblot was ealized with Anti-P53 antibody.

## Results

### P53 -fused to GST-decreases bacterial growth preferentially at low temperature

The GST-P53 fusion protein shows partial inhibition of recombinant strain BL21(DE3), **Fig 1**. This is observed when bacteria are induced early -from the start-in growth kinetics with IPTG concentrations usually used in purification experiments (0.2 mM here). The growth inhibition is well detected from the exponential phase of bacterial cultures and continues throughout the stationary and decline phases. This growth attenuation is mainly observed at low temperatures (25°C) (and 30°C: data not shown) rather than at 37°C.

**Fig 1:**
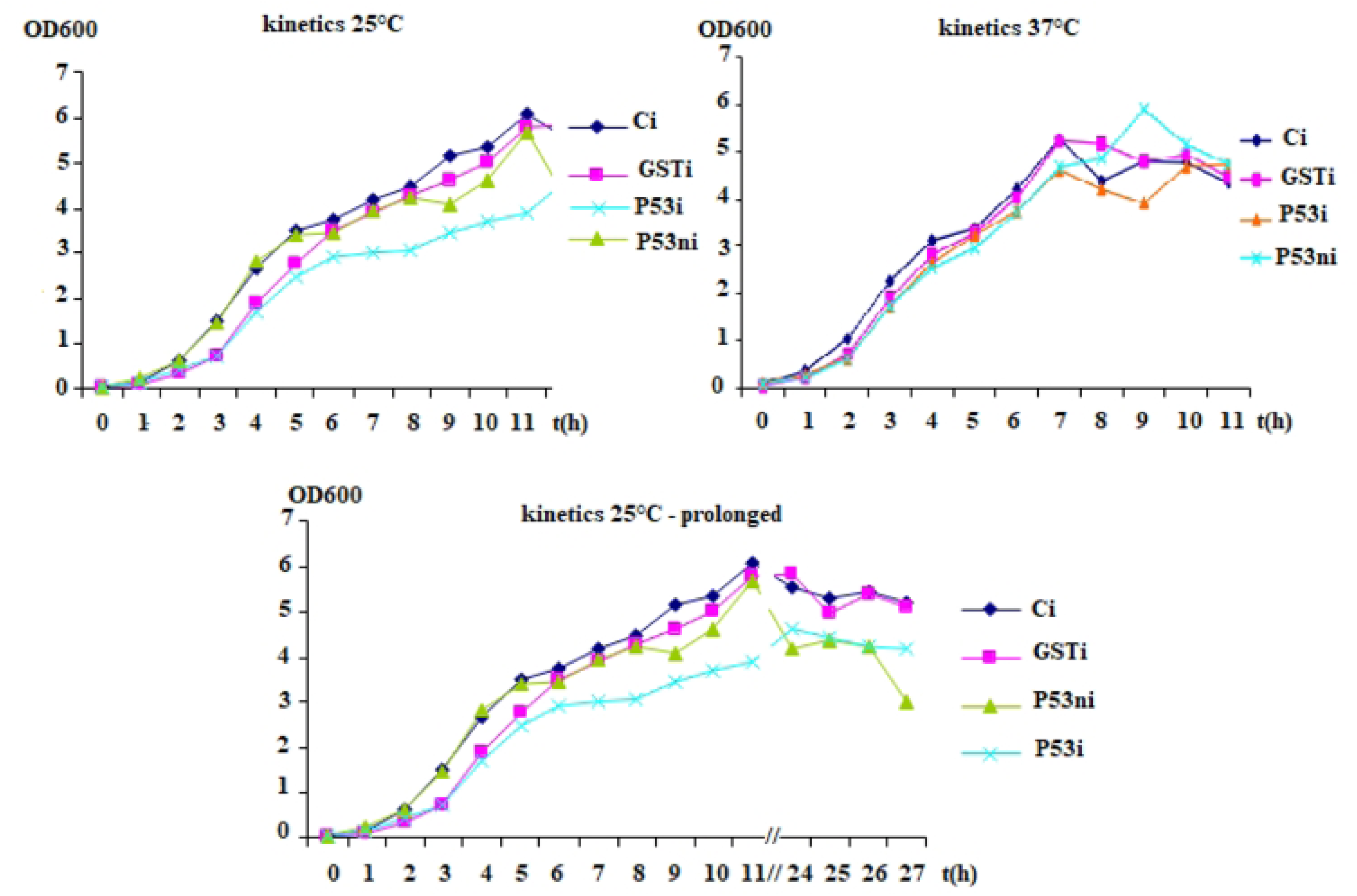
Growth kinetics of BL21 (DE3) E. coli strain expressing GST-P53. Three strains are used for growth kinetics at two temperatures 25°C and 37°C in four conditions: Ci, Control non recombinant strain, GSTi, GST-expressing pGEX4T3 harboring strain, P53i, pGEX4T3-P53 recombinant strain, all three induced and P53, non induced pGEX4T3-P53 recombinant strain. Induction was performed at the start of the cultures with 0.2 mM IPTG. Same number of cells are assumed for the inoculum by Malassez counting. a) 25°C kinetics b) 37°C kinetics and c) 25°C kinetics pursued one night.

The effects observed with non-induced recombinant clones can be atrributed to an expression leakage existing with the pGEX system. The non-induced strain expressing the GST-P53 fusion eventually acquires a delayed growth curve after 10-11 hours of kinetics at 25°C. The decline for this non-induced GST-P53 strain at the end of kinetics seems to be more marked than for the induced strain or for the two other control strains, the non recombinant (NR) and the GST.

The GST moiety shows itself a degree of growth inhibition that is particularly noticeable at the beginning of the exponential phase at 25°C. However after this phase and in general, it remains closer to the non-recombinant (NR) strain than to the P53 fusion.

In other low temperature kinetics (25°C and 30°C), probably with a different concentration of inoculum, a total inhibition of induced GST-P53 strain could even be observed up to 7-11h, compared to a greater growth for the non induced form, or the induced control or the GST strains (data not shown).

### Variability between strains : BL21(DE3) produces very high expression resistant to post-lysis degradation

BL21(DE3) was initially chosen to express P53 because it has a good expression level as shown by the western blot, **Fig 2**. Moreover, compared to other strains (DH5α, TG1, TOP10 and XL1Blue) by the same immunoblotting assay, the expression level of BL21(DE3) is the highest under the same conditions. The strain also confers some resistance to post-sonication degradation unlike other strains.

**Fig 2:**
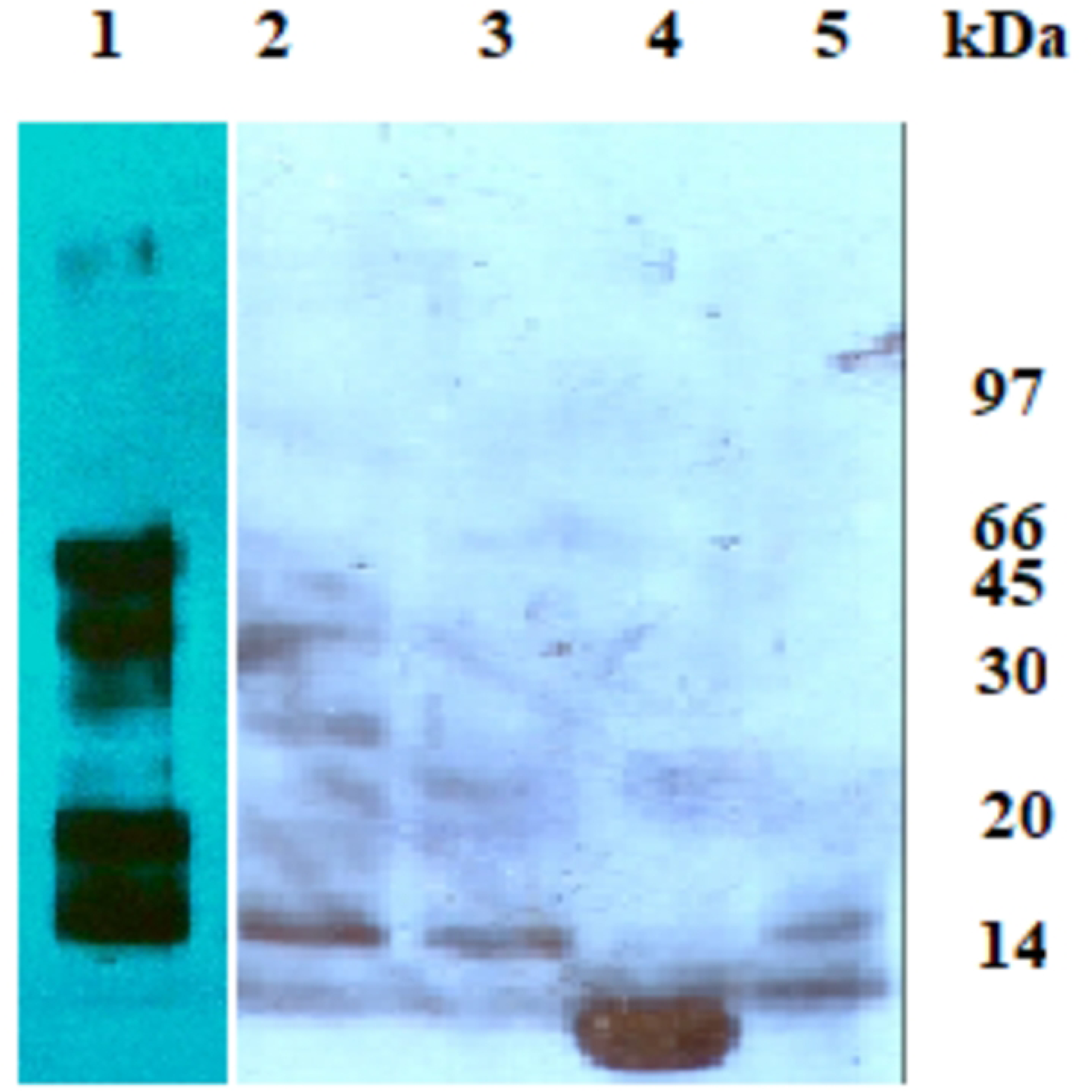
P53 expression in lysates of bacteria strains revealed by Western blot. Five strains BL21, Dh5α, TG1, Top10 and Xl1Blue expressing GST-P53 are cultivated at 37°C till OD_600_ 0.6-0.8, induced with 0.5Mm IPTG for 3 h at 30°C. Cells are lyzed with sonication (in MOPS buffer, PMSF and Lysozyme). After migration and transfer, blots were revealed with anti-P53 antibody. 1-5 : lysates of BL21, Dh5α, TG1, Top10 and Xl1Blue, respectively. We can even suspect that sonication has cleaved between GST and P53 since the size of the fusion should be greater (around ∼75kDA).

Another aspect related to the high expression of P53 by BL21(DE3) is the propencity to protein degradation or truncation, as shown by Western blot results. This may be related to the overproduction or an aggregation or something else. P53 degradation/truncation is less noticed in the other strains, or at least less visible because there are fewer protein. In addition, the BL21 profile also shows the existence of oligomers or large compounds. **Fig 2**

### Addition of Ampicillin to an induced GST-P53 culture causes transient inhibition

Ampicillin, usually pre-added to the culture media of recombinant strains, was introduced once in the GST and GST-P53 cultures in the middle of the exponential phase (around 4:30 hours) and not at the beginning of culture, following a fortuitous omission. The cultures were carried out at 25°C and previously induced by IPTG from the beginning. This caused a rapid drop in OD600 and the appearance of viscosity with the GST-P53 culture but not with GST. And these signs continued for a few hours before resuming growth with a rise in OD600, **Fig 3a**.

**Fig 3:**
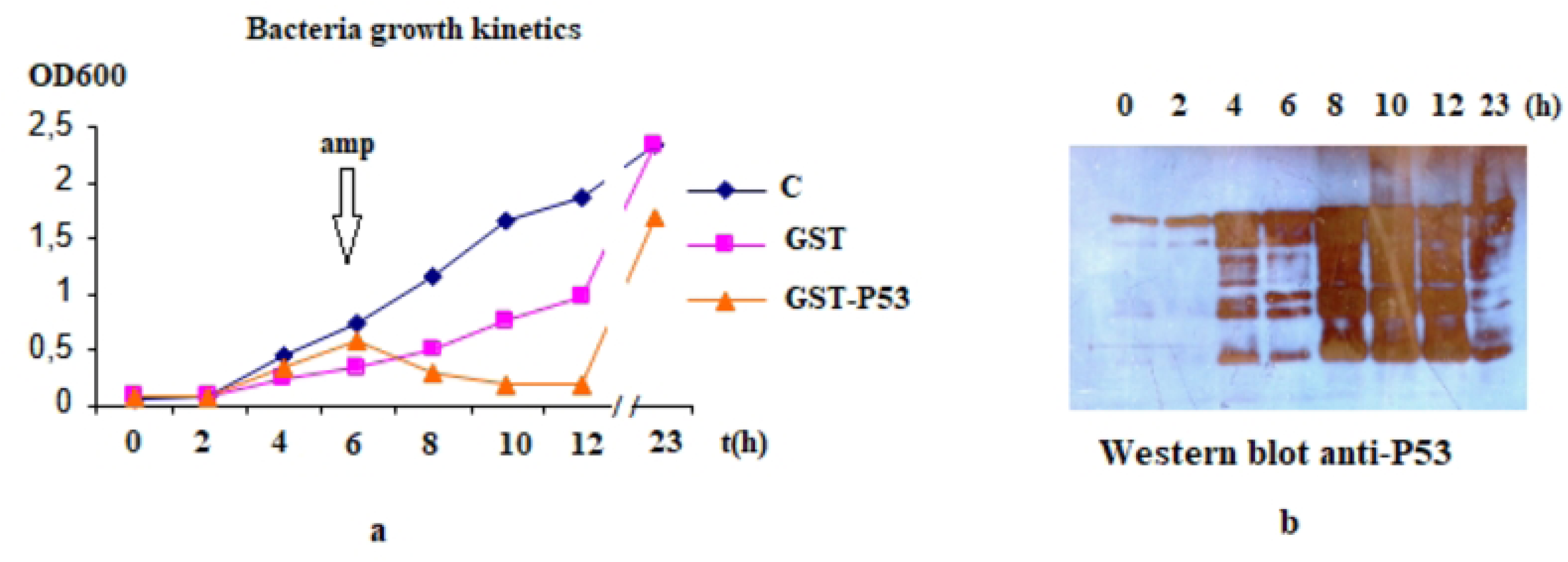
Effect of ampicillin supplementation in the middle of induced P53-expressing bacterial culture. a) Growth kinetics. Three BL21(DE3) bacteria strains : C, non transformed, GST and GST-P53 expressing respectively GST and the fusion GST-P53 ; were grown at 25°C. The two transformants were induced with IPTG from the beginning. Ampicillin was not added to the medium beforehand but was introduced in the middle of the kinetics (exponential phase) in the culture medium of the two recombinant strains at the usual concentration (100 µg/ml). This kinetics experiment was repeated once more with similar results of optical density drop following the introduction of ampicillin. b) Monitoring of P53 expression by Western blot during culture.

The previous kinetics with ampicillin added from the beginning **(Fig1)** is considered as a control for this experiment.

Culture specimen corresponding to the OD_600_ sampling were migrated in an SDS-gel and the P53 revealed by Western blotting. Interestingly, the P53 protein signal increased sharply directly after the addition of ampicillin. And this remained so for a few points until the p53 signal was recovered more moderate. The imunoblot profile corresponds exactly to that of growth kinetic. The increase in the P53 signal coincides with the decrease of OD_600_ and the return to lower P53 level coincides with an ascent of the same OD_600_ **Fig 3b**. Once the antibiotic is taken up by the cells and its effect has faded after some time (more than 4-5 hours and perhaps overnight), the P53 protein signal decreases and becomes moderate again and OD_600_ is high again as recorded one night after the start of culture. The subsequent OD_600_ always remains lower than that of GST and non recombinant strains. Concerning the strain expressing GST, which was subjected to the same treatment, i.e culture on non-antibiotic medium then introduction of ampicillin in exponential phase, the inhibitory effect could be observed throughout the kinetics compared to the control (non-recombinant BL21 strain), more pronouced than the previous one. However, there was no massive drop in optical density as with GST-P53.

### Persistant P53 in a test of phage incubation in P53-expressing bacteria

In a previous work, a phage display library screen yielded a phage-peptide clone recognizing GST-P53 fusion protein **[2]**. The phage clone was named Ph6. An experiment was performed as follows : *E. coli* strain BL21(DE3) expressing GST-P53 (previously induced) was infected separately with three phage clones including Ph6. The other phages were : M13mp18 and Ph17 (a non-recombinant empty phage from phage display libraries), both as controls. After overnight culture, the phage-infected bacteria were viscous with foam indicating cell lysis (data not shown). And the culture-infection tubes were left (at 4°C) for long months. An SDS-denaturing coomassie blue gel was performed with all total bacterial cultures in this assay. The Ph6-infected strain showed a distinctive profile as shown in **Fig 4.a**. While some bands disappeared compared to the other conditions, others can clearly be visible.

**Fig 4:**
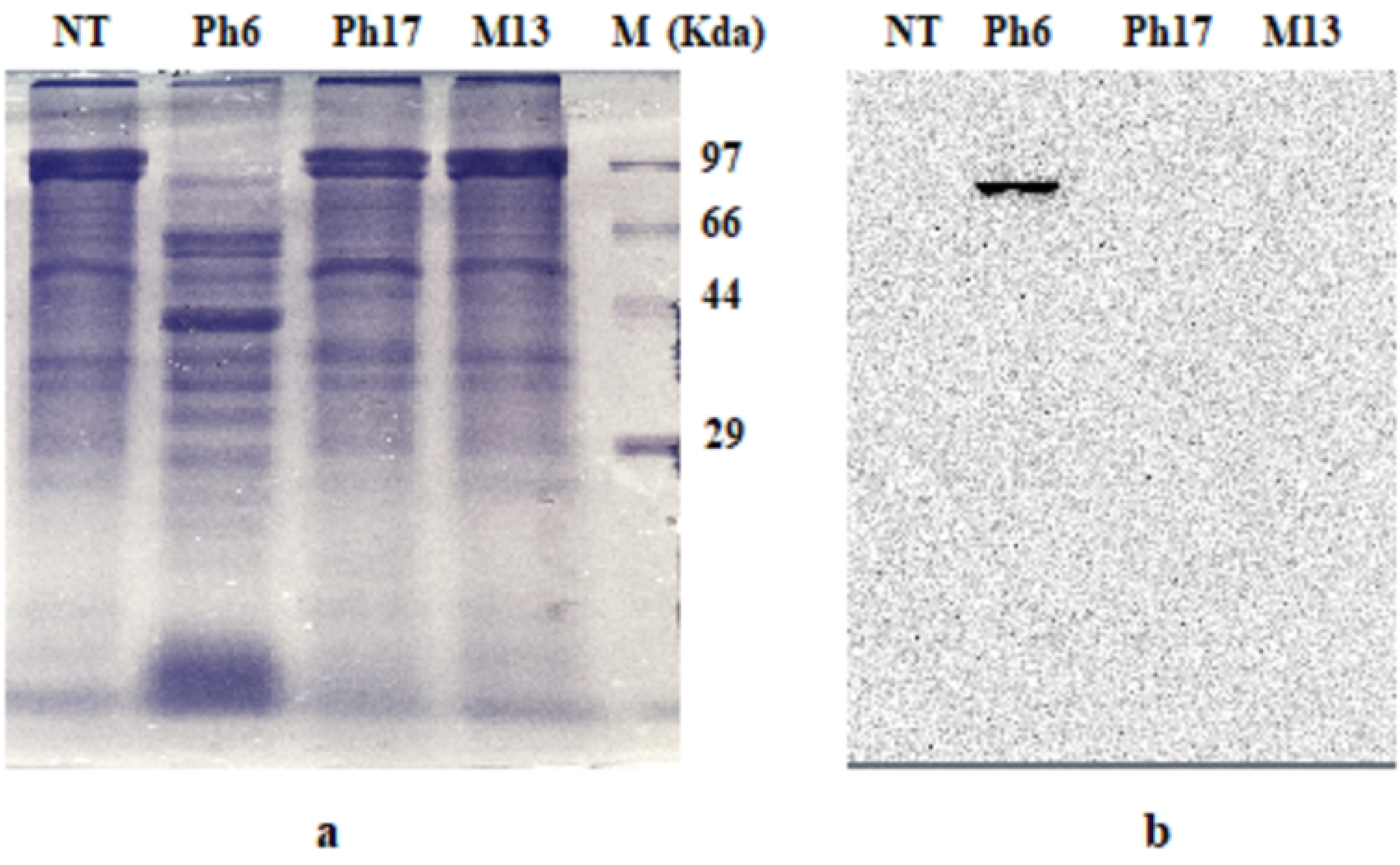
SDS-denatured protein gel and immunoblot of BL21 GST-P53 infected with distinct phages. Induced cultures of BL21 GST-P53 strain were infected separately in early exponential phase with three phages : Ph6, a P53-specific phage, Ph17, an empty phage and M13 : M13mp18 ; with an untreated control (NT). The infection was left overnight and cultures were migrated later (a). Immuno-blot was visualized with Versadoc (b).

The most relevant result came from the immunoblot assay which showed a good GST-P53 fusion signal, abundant in one band of ∼75kDa in the lysed slimy culture of the P53-expressing strain infected with Ph6. All other cultures (Ph17 an empty phage, M13mp18, non infected) did not show similar signal. P53 is either partially or totally degraded in these cultures **Fig 4.b**. Another remark is that GST-P53 was found in good quantity and quality (non degraded) many months after the culture tubes were stored at 4°C And this despite a cell lysis evident to the naked eye.

## Discussion

BL21(DE3) produces a very high expression but this is not necessarily positive. This high level of GST-P53 expression by BL21(DE3) can be a challenge for the physiological aspect, mainly cell growth. In addition to the conventional 37°C temperature, (25-30°C) were tested. Exactly, a growth inhibition becomes evident under low temperature conditions. Considering the possible parallel with the protein effect in yeast, a milder P53 expression/dosage results in milder growth inhibition in *Pichia pastoris* **[8]**. It can be assumed that the same can be found in bacteriaP53 overexpression is likely more inhibitory than moderate/low expression. BL21(DE3) can therefore be considered as the overexpression mode here in the bacterial context. This is demonstrated by the expression levels as shown by immunoblots of distinct strains. It seems that *E.coli* physiology and growth may also be affected by this P53 overexpression too.

This finding supports the idea of a functional P53 when expressed in *E. coli*. The protein probably escaping aggregation, adduction and insolubility by low temperatures, recovers some functions. Reversing the aggregation of the P53 protein restores its functionality **[18]** and thus its effect on cell growth in this case. As it may have simply gained functions.Another hypothesis can be made on a temperature degrees preference of about 25-30°C for the heterologous P53 protein to be active and functional in bacteria as in yeast ; independently of the question of solubility. The culture temperature of yeast is 30°C. This temperature preference may be a common condition between the two organisms.

The actions of P53 are generally triggered by the induction of stress ; and this generally indirectly, via variable effectors **[19]**. We obtained an inhibitory effect of the introduction of ampicillin into an already induced GST-P53 BL21 culture. We conclude from this test that the addition of ampicillin triggered a reaction of cells already expressing certain quantities of GST-P53 since induced from the beginning of the culture. Cells that were besides, grown and induced at (25°C), temperature which allowed the inhibitory effect in normal kinetics before. Ampicillin can be detected/considered as a stress applied to bacteria and this would haver led to an accumulation of P53 protein. The latter caused a fall of OD_600_ reflecting with viscosity a cell lysis and a decrease in growth.

This inhibition was transient and growth resumed after a few hours. It is possible that induction became weak, that P53 was no longer synthesized and that the residual protein lost its effect. We can as well think of an emergence of mutants/revertants without or with a lesser inhibitory effect, as previously described for the yeast *Saccharomyces cerevisiae* **[20]**.

We found that already in 2012, researchers described an apoptotic effect of antibiotic treatments of *E. Coli* with apoptosis hallmarks and that RecA was identified as the partner that binds the peptide substrates of eukaryotic caspases **[21]**.

The phage infection test of P53-expressing bacteria showed that there could be a relationship (association or interaction) between the expression/availability of the P53 protein in *E.coli* and phage infection as well as with particular infection with a phage recognizing P53.

Our hypotheses to explain the cell lysis of infected bacteria expressing GST-P53: either P53 caused a reaction to infection by phage as if it were an aggression or stress and caused cell death processes (probably an apoptosis). Or it was the phage itself that caused the cell lysis due to the presence of P53 and/or helped by it. This despite the fact that BL21(DE3) is a F-strain, therefore theoretically incapable of being infected by the phage because it does not have cellular pili. However the long-term incubation may have overcome pili infection.

Interestingly, this is reminiscent of abortive infection, a bacterial immune mechanism that involves, among other processes, bacterially regulated cell death to save an entire population **[22]**.

The fact that only Ph6, which is a phage presenting a heptapeptide recognizing P53 (NPNSAQG) **[2]**, produces this effect : persistence and integrity of the P53 protein, reflects a possible precise role of this specific peptide. The fact that the P53 protein is still present and intact after weeks of preservation of the phage-infected culture is also a curious fact. Did an interaction/binding between the protein and its phage-peptide ligand protect it (or both) from degradation ?

The test and its surprising results suggest the idea of a potential use of phage infection and proteins specific phage infection as tools for bacterial cell lysis. The use of phages is not a new method, it is known as phage therapy **[23]**. But this specificity in targeting expressed proteins or precise molecules is of some interest.

Recall that P53 orthologs exist in some eukaryotes but not in yeast or prokaryotes (bacteria) so far. Recently, however, a protein was discovered in the yeast *Saccharomyces cerevisiae* with DNA Binding homology to p53 **[24]**. And more lately, a tetramerization domain has been found in prokaryotic and eukaryotic transcription regulators that is homologous to p53 **[25]**. This strengthens our theory that there is an active or acting p53 or p53-like factor in prokaryotes. Possible interference with GST existing either as fusion fragment or as a separate protein obtained from a truncation is also possible too, especially for the effect on growth.

## Conclusion

Bacterial growth kinetics and a number of other physiological observations suggested to us some roles for P53 when expressed in Escherichia coli (mainly the strain overexpressing BL21(DE3) and under certain temperature conditions below 37°C). This includes cell growth and attenuation of cell viability which can be any of these processes: control/arrest of cell growth, cell death, cell lysis, etc. Cell lysis - death - was the ultimate fate of this bacterium when infected with phages. A special reaction was observed when P53 and a specific phage peptide were present simultaneously in the bacterial culture. This led us to speculate that the recombinant human P53 protein expressed in bacteria might be active at the cellular physiological level. And that the protein might possibly act in a manner similar to that expressed in some cell lines and yeast (eukaryotes); it could trigger stress reactions, order and execute the arrest or inhibition of cell growth up to cell death. Particular attention must still be paid to inter-strain variabilities.

## Acknowledgments

We thank the current and former members of the Laboratoire de Biotechnologie Moléculaire des Eucaryotes (LBME) who helped us in this work.

We ascertain that no AI tools were used to produce this manuscript.

All data are made available at 10.6084/m9.figshare.26490085.

